# Atomic force microscopy reveals new biomarkers for monitoring subcellular changes in oxidative injury: neuroprotective effects of quercetin at the nanoscale

**DOI:** 10.1101/353557

**Authors:** Maja Jazvinšćak Jembrek, Josipa Vlainić, Vida Čadež, Suzana Šegota

**Affiliations:** Division of Molecular Medicine, Ruđer Bošković Institute, Bijenička cesta 54, Zagreb, Croatia; Department of Psychology, Croatian Catholic University, Ilica 242, Zagreb, Croatia; Division of Physical Chemistry, Ruđer Bošković Institute, Bijenička cesta 54, Zagreb, Croatia

**Keywords:** quercetin, Akt and ERK signalling, atomic force microscopy, roughness, Young’s modulus, elasticity

## Abstract

Oxidative stress is a process involved in the pathogenesis of many diseases, including atherosclerosis, hypertension, diabetes, Alzheimer’s disease etc. The biomarkers for assessing the degree of oxidative stress have been attracting much interest because of their potential clinical relevance in understanding cellular effects of free radicals and evaluation of the efficacy of drug treatment. Here, an interdisciplinary approach using atomic force microscopy (AFM) and cellular and biological molecular methods were used to obtain new potential biomarkers for monitoring oxidative stress condition. Biological methods confirmed the oxidative damage of investigated P19 neurons and revealed the underlying mechanism of quercetin protective action. AFM was employed to evaluate morphological (roughness) and nanomechanical (elasticity) properties that may be specific biomarkers for oxidative stress-induced cytoskeletal reorganization manifested by changes in the lateral dimension and height of neuronal somas. The morphological and nanomechanical analysis of neurons showed the strong mutual correlation between changes in cell membrane elasticity and neuroprotective effects of quercetin. Our findings indicate that AFM is a highly valuable tool for biomedical applications, detection and clarifying of drug-induced changes at the nanoscale and emphasize the potential of AFM approach in the development of novel therapeutic strategies directed against oxidative stress-induced neurodegeneration.

## Introduction

Oxidative stress, which occurs when the cellular antioxidant defence is insufficient to keep the levels of reactive oxygen species (ROS) below a toxic threshold, represents a common mechanism of neuronal death in a variety of neuropathologies, including neurodegenerative diseases such as Alzheimer’s disease and Parkinson disease [1]. The brain is particularly vulnerable to oxidative stress due to its large oxygen consumption, high content of polyunsaturated fatty acids, accumulation of redox-reactive transient metal ions and limited endogenous antioxidant protection [2].

At physiological levels ROS act as signalling molecules, but when present in excess they may induce an oxidative stress response and trigger cell death by modulating redox-sensitive signalling pathways and gene expression. Among different ROS molecules, H_2_O_2_ is considered as the key target in neuroprotection as is one of the most abundant ROS in aerobic organisms. Moreover, it can be converted to more toxic species of which hydroxyl radical is particularly dangerous [3]. The mechanism of the H_2_O_2_-mediated signalling relies on the oxidation of redox-sensitive thiol groups in cysteine residues of different target enzymes and transcription factors, thereby modulating their functions [4]. At concentrations above the physiological threshold, H_2_O_2_ can change activities of different signalling cascades, such as pathways of mitogen-activated protein kinases (MAPK) and protein kinase B (PKB/Akt). This ultimately triggers specific nuclear or cytoplasmic response, often ending in cell death [5,6]. In addition to modulation of intracellular transduction pathways, increased levels of ROS may induce damage to all biological macromolecules that further exacerbates neuronal death [7].

Quercetin, a plant-derived polyphenolic nutraceutical, possesses a wide spectrum of health-promoting effects mainly attributed to its strong antioxidative capacity. Both *in vitro* and *in vivo* quercetin is effective against various oxidants and other neurotoxic molecules that induce oxidative stress and mimic the pathological hallmarks of neurodegenerative diseases [8,9]. *In vitro,* quercetin exerts its neuroprotective effects acting as a potent direct radical scavenger and a metal chelator but is also able to downregulate redox-sensitive signalling [10,11]. Due to low bioavailability, the expected concentrations of quercetin in the brain tissue are below those that are required for direct antioxidant activity. Furthermore, concentrations of endogenous antioxidants greatly surpass levels of quercetin in the brain. Hence, it is suggested that the neuroprotective effects of quercetin *in vivo* are not achieved through direct ROS scavenging.[9] Instead, it seems that the modulation of intracellular signalling pathways could represent a primary mode of quercetin action *in vivo* [12,13]. In support of this assumption, it is shown that structural characteristics of quercetin molecule involved in neuroprotective action differ from those that provide free radical scavenging [14].

Inside the cell quercetin accumulates in the nucleus, at distinct loci, where may affect transcription and/or activity of numerous transcription factors [15]. For example, induction of NF-E2-related factor-2 (Nrf2) transcription factor pathway, which drives the expression of antioxidant genes and increases endogenous antioxidant defence, is one of the well-recognized mechanisms of quercetin action that greatly contributes to neuroprotection [9,16]. Quercetin also may act inhibitory on a number of kinases signalling pathways including Akt/PKB, extracellular signal-regulated protein kinases (ERK) 1/2 and c-Jun N-terminal kinase (JNK) [12,17,18]. The stimulatory effects on the same kinases are also demonstrated, leading to the expression of survival and defensive genes [11,19]. Hence, the exact mechanism of neuroprotective effects of quercetin remains puzzling, particularly when considering modulation of intracellular signalling pathways.

Physiological functions of a cell are closely related to its morphological characteristics [20,21]. When pathological toxin-and drug-induced molecular changes occur inside the cell, its overall morphology usually changes as well. Recent studies have shown that the nanomechanical behaviour of the cell plays an important role in maintaining cellular physiological functions and also presents a novel biomarker for indicating different cell states [22]. In the present study, we implemented a novel experimental approach, combining molecular biology with atomic force microscopy (AFM) as an advanced tool, to obtain information about drug-induced neuronal changes during oxidative stress. The field of AFM applications in neuronal research is developing. AFM is primarily used for quantitative imaging of surface topography. In non-imaging mode, AFM provides spatially resolved maps of the nanomechanical characteristics usually reported as cell elasticity [23,24]. Both imaging and non-imaging AFM modes can bring specific information about neuronal membranes and cytoskeleton architecture. Structural and nanomechanical properties of neurons are highly affected by environmental conditions that can change the organization of their microtubules, actin filaments and neurofilaments [24,25]. AFM was already employed for detecting nanoscopic changes resulting from oxidative damage in the plasma membrane of glioblastoma cells [26].

The aim of the present study was to combine capabilities of AFM with molecular biology tools to better understand cellular and molecular consequences of H_2_O_2_-induced oxidative injury in P19 neurons and to investigate molecular mechanisms of quercetin-mediated neuroprotection that falls beyond its antioxidant activity. For the first time, we used AFM to reveal subtle changes of neuronal membrane topography and nanomechanical properties induced by drug treatment during the oxidative insult.

## Materials and methods

### Chemicals and Reagents

*all-trans* retinoic acid (ATRA), cytosine-arabinofuranoside, poly-L-lysine, 1,4-diamino-2,3-dicyano-1,4-bis[2-aminophenylthio] butadiene (UO126) and all chemicals used for maintaining and differentiation of P19 cells (unless otherwise stated) were purchased from Sigma-Aldrich Chemicals (St. Louis, MO, USA). Quercetin dihydrate was obtained from Aldrich Ch. Co. Inc. (Milwaukee WI, USA), wortmannin was purchased from Ascent Scientific (Princeton, NJ, USA) and hydrogen peroxide solutions were obtained from Kemika (Zagreb, Croatia) and Sigma-Aldrich Chemicals (Cat. No. 216763, stabilized solution containing inhibitors). All other chemicals used were of analytical grade.

### P19 cell culturing and P19 neuronal differentiation

Undifferentiated P19 cells (pluripotent mouse teratocarcinoma cell line) were cultured in high-glucose Dulbecco’s modified Eagle’s medium (DMEM) containing 10% heat-inactivated fetal bovine serum (FBS), 2 mM L-glutamine, 100 units/ml penicillin G and 100 μg/ml streptomycin (growth medium) in a humidified atmosphere of 5% CO_2_ at 37°C. They were passaged every two days using trypsin (0.05% trypsin, 1 mM EDTA) in phosphate buffered saline (PBS).

For induction of neuronal differentiation, exponentially growing P19 cells (1×10^6^) were seeded into nonadhesive bacteriological-grade Petri dishes (10 cm) containing 10 ml of DMEM medium supplemented with 5% FBS, 2 mM L-glutamine, antibiotics and 1μM ATRA (induction medium). Embryonal bodies of P19 cells were formed in 1-2 days. After 48 h, the old medium was replaced with the fresh ATRA-containing medium and aggregated cultures were grown for two more days. After the four-day of ATRA treatment, P19 embryonal bodies were harvested, washed with PBS, trypsinized, collected by centrifugation (200 g, 5 minutes), and resuspended in growth medium. For optimal neuronal differentiation single cells at a density of 10^5^ cells/cm^2^ were plated onto 96-well plates or 35 mm Petri culture dishes (Cell^+^, Sarstedt, Newton, NC, USA and NUNC, Roskilde, Denmark), and grown in growth medium for two more days. Finally, the growth medium was replaced with serum-free medium containing DMEM supplemented with insulin, transferrin, selenium and ethanolamine solution (ITS-X, Gibco), 2 mM L-glutamine and antibiotics (neuron-specific medium), and cells were grown for additional 2 days in the presence of 10 μM mitotic inhibitor cytosine-arabinofuranoside (AraC) to inhibit proliferation of non-neuronal cells. Complete neuronal maturation was confirmed with monoclonal anti-tubulin β-III mouse IgG, clone TU-20 conjugated with Alexa Fluor^®^488 (Millipore, Temecula, CA). Differentiated cells expressing neuronal marker β-III tubulin were visualized by fluorescence microscopy (data not shown). Completely differentiated P19 neurons exert morphological, neurochemical and electrophysiological properties resembling neurons from the mammalian brain, and represent an established model for pharmacological studies.[27,28]

### Drug treatment

In all experiments, P19 neurons were treated 8 days after the initiation of differentiation procedure (DIV8). Each batch of cultured cells was divided into control and drug-treated groups. For inducing oxidative damage, P19 neurons were incubated with 1.5 mM H_2_O_2_ in the neuron-specific medium for 24 hours, alone or in the presence of various concentrations of quercetin that failed to affect the viability of P19 neurons when applied alone.[28] To examine the effects of quercetin on kinase signalling pathways, P19 neurons were pretreated with UO126 or wortmannin for 60 min and then exposed to H_2_O_2_, inhibitor and 150 μM quercetin for additional 24 hours.

## Assessment of cell death

### Trypan blue exclusion assay

The viability of P19 neurons in the presence of 1.5 mM H_2_O_2_ was analysed by trypan blue exclusion assay. The method is based on the principle that healthy cells effectively exclude the dye from their cytoplasm, while those with damaged membranes lose this ability and appear blue. Following treatment, culture mediums with floating cells were collected in a centrifuge tube. Attached cells were trypsinized for 5 min, resuspended and pooled with the corresponding medium. Samples were centrifuged at 250 g for 5 min; pellets were resuspended in 250 μl of neuron-specific medium and incubated for 5 min in the presence of 0.4% trypan blue solution. The ratio of trypan blue stained nuclei over the total number of cells was used to determine the percentage of cell death. For each examined group at least 500 neurons were counted by two different investigators.

### MTT assay

Effects of inhibitors of Akt and ERK1/2 signalling, wortmannin and UO126, respectively, on the neuroprotective effect of quercetin, were determined by 3-(4,5-dimethylthiazol-2-yl)-2,5-diphenyl tetrazolium bromide (MTT) assay. MTT assay was also employed to determine effects of different concentrations of H_2_O_2_ on the viability of P19 neurons. Estimation of viability is based on the ability of P19 neurons to cleave dissolved MTT into an insoluble formazan product by cleavage of the tetrazolium ring by dehydrogenase enzymes. Briefly, P19 neurons were seeded on 96-well micro-plates and were incubated for 24 hours with H_2_O_2_, quercetin and inhibitors. At the end of treatment schedule, the medium was removed, and the cells were incubated with 40 μl of MTT solution (0.5 mg/ml final) for 3 h at 37°C. Precipitated formazan was dissolved by adding 160 μl of dimethyl sulfoxide (DMSO). Optical densities of coloured solutions in each well were determined by an automatic ELISA reader at 570 nm. The data were analysed after blank subtraction from all absorbance readings and viability of P19 neurons was calculated according to the following equitation: % cytotoxicity = [A_control_-A_treatment_]/A_control_ x 100.

### Measurement of reduced glutathione

Reduced glutathione (GSH) is one of the major non-enzymatic intracellular antioxidant defence mechanisms. Changes in the intracellular level of GSH were monitored by using a GSH-Glo^TM^ Glutathione Assay (Promega, Madison, WI, USA) based on the conversion of a luciferin derivative into luciferin by glutathione S-transferase (GST) in the presence of GSH. Hence, the signal generated in a coupled reaction with luciferase is proportional to the amount of GSH present. According to the manufacturer’s instruction, following treatment, the medium was removed and 100 μl of GSH-Glo^TM^ reagent was added per well. After a 30-minute incubation, 100 μl of luciferin detection reagent was further added and following 15-minute incubation emitted light was measured in the luminometer (Fluoroskan Ascent FL, Thermo Scientific).

### Determination of Bcl-2, Bax, p53 and GAPDH mRNA levels by semi-quantitative RT-PCR

Expressions of Bcl-2, Bax, p53 and GAPDH mRNA were examined by semiquantitative RT-PCR analysis according to the method previously described by Jazvinšćak Jembrek and co-workers.[28] cDNAs were amplified and analysed during two consecutive cycles in the log phase of PCR reactions. PCR primers, annealing temperatures, and numbers of cycles are shown in Table 1. The reactions were performed in a Perkin Elmer 9600 thermocycler. Amplified products (10 μl) were electrophoretically separated on a 1.5% agarose gel and stained with ethidium bromide (0.5μg/ml) for 20 minutes. Optical densities of detected bands were analysed using ImageJ NIH software 1.0. Expression of housekeeping gene TATA-binding box protein (TBP) mRNA was used as an internal standard for normalization.

**Table 1.**
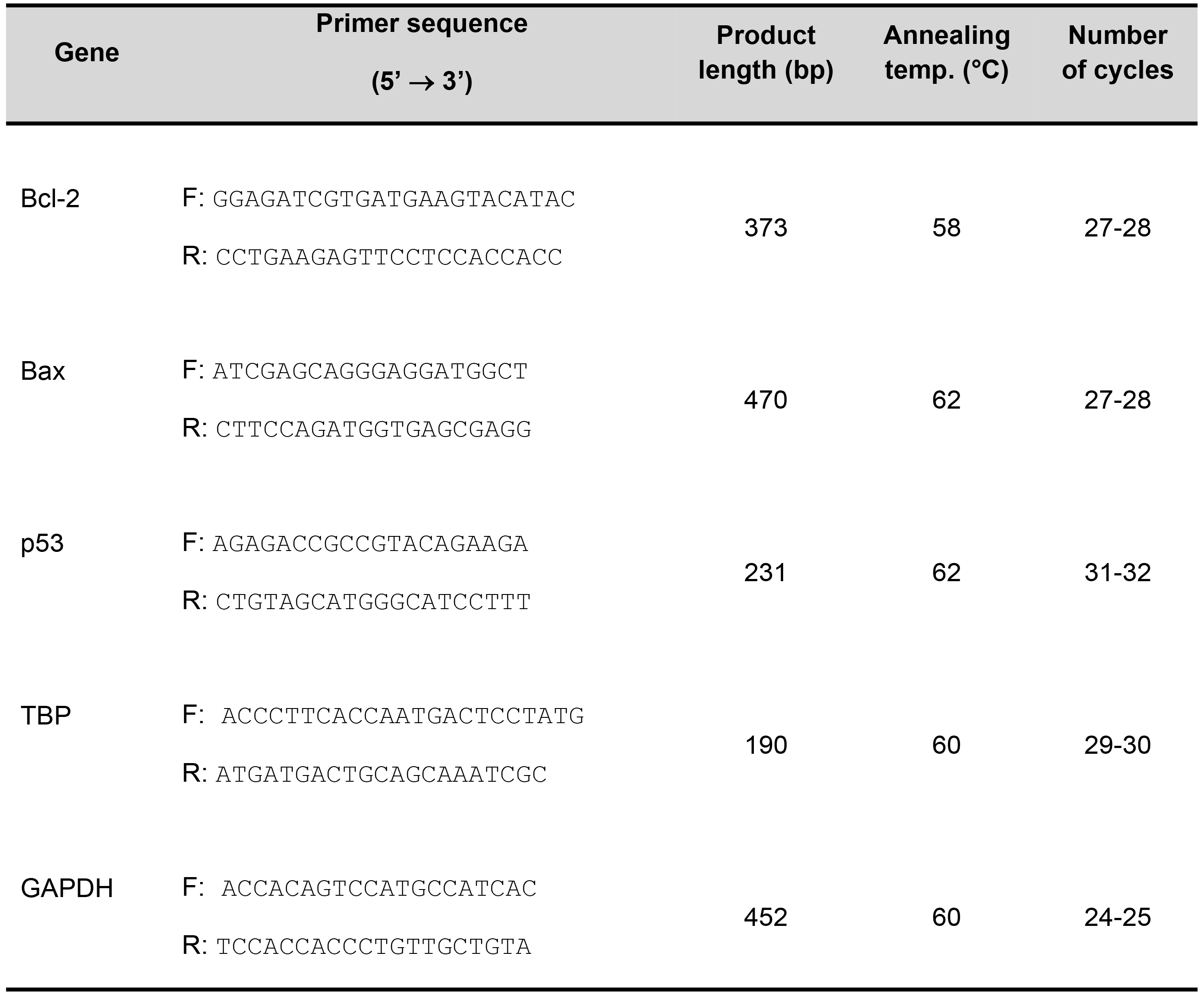
Primer sequences and conditions used for PCR amplifications

### AFM measurements

All cell imaging and force mapping measurements were obtained on the JPK NanoWizard^®^ ULTRA Speed and JPK NanoWizard^®^ 4 AFM system coupled to the Nikon Eclipse TE2000-U inverted optical microscope. Cantilever qp-Bio-AC with the nominal spring constant of 0,05-0,07 N/m was used for measuring in liquid. The measurements are performed in PBS with the Petri dish fixed to the standard sample holder. All images were acquired from fixed neurons to provide a more accurate measure of the height and structure of the individual subcellular regions using QI^™^ mode. Neurons grown on Petri dishes were fixed with 4% paraformaldehyde (for 15 minutes at RT). The Petri dishes with the sample cells were fixed on the standard sample holder using two-component rubber glue. Before imaging, neurons were examined with an inverted microscope, fresh PBS was added, and an interesting cell was selected. The focus was on neurons with a round cell body. Furthermore, morphological properties of the distinct soma regions were determined at the optimized imaging conditions to avoid any possible influence of the applied force on imaging. During scanning both trace and retrace images were recorded and compared for accuracy. No substantial difference could be observed between them. For each sample, an overview scan of the whole cell was acquired, as well as a detailed image of 2.5 μm x 2.5 μm with 256 μ 256-pixel resolution. For all measurements, a setpoint of 400 pN and an extend/retract speed of 110-195 μm/s was used. For the overview image of the control sample, a setpoint of 500 pN was used. Prior to measuring, the cantilever was calibrated using the in-built non-contact method.[29] The QI™ data were analysed using the JPK Data Processing software.

The roughness of the detailed images was calculated for the raw height image and for the height image treated with a quadratic plane fit in order to minimize the influence of the cell surface curvature on the roughness values. The plane fit calculates a plane of the desired order from the raw image and subtracts this plane from the whole image. Roughness Average, Ra, is the arithmetic average of the absolute values of the profile heights over the evaluation length. It is defined as

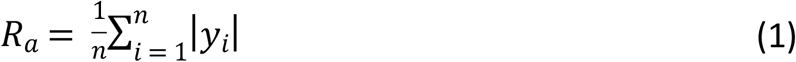

where *y*_i_ are vertical deviations height profiles from the mean plane, while RMS Roughness, *R*_q_, is the root mean square average of the profile heights over the evaluation length defined as:

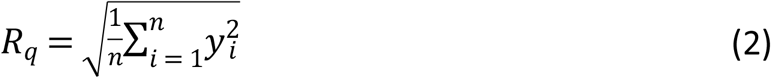

The Young’s modulus was determined using the Hertz model for conical indenters.

### Statistical analysis

Statistical analysis of the data was carried out using GraphPad Software (San Diego, CA). All values are represented as mean ± SEM from at least three independent experiments. Comparisons between group means were evaluated by one-way ANOVA, and when statistically significant, post hoc analysis with Dunnett’s multiple comparison test or Tukey’s test followed. The P values less than 0.05 were considered statistically significant.

## Results and Discussion

### Biological approach reveals neuroprotective effects of quercetin in H_2_O_2_ induced injury

Exposure to H_2_O_2_ in a concentration-dependent manner decreased the viability of P19 neurons (Figure 1A). P19 neurons tolerated relatively high concentrations of quercetin, up to 150 μM (Figure 1B). Quercetin applied together with 1.5 mM H_2_O_2_ improved survival of P19 neurons indicating the neuroprotective effect of quercetin against oxidative injury. Quercetin action was dose-dependent, and at the highest concentration applied it maintained viability at 87.6 ± 6.7 % of the control group (Figure 1C).

**Fig 1.**
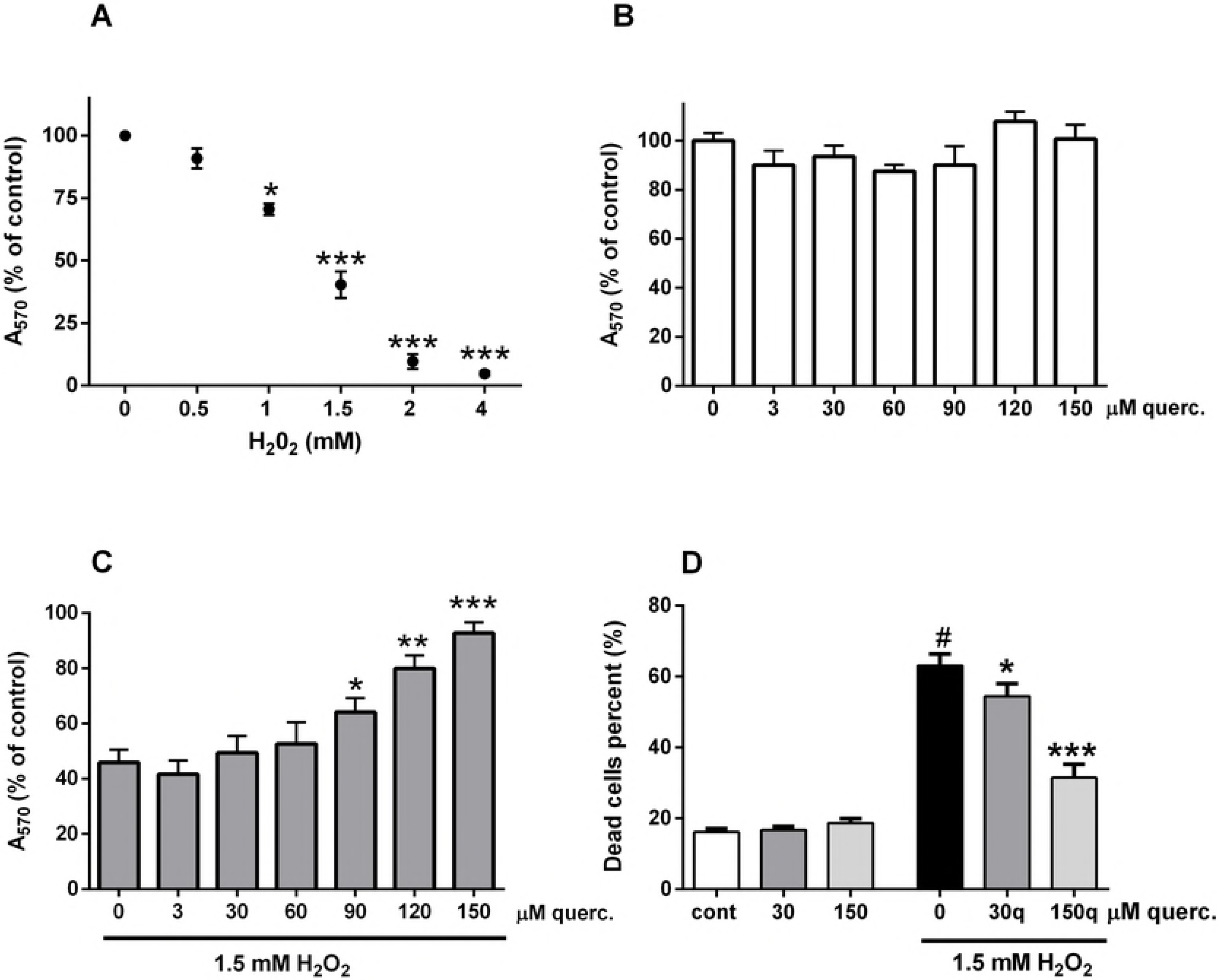
Quercetin improved viability of P19 neurons exposed to H_2_O_2_. In a dose-dependent manner, H_2_O_2_ reduced the viability of P19 neurons (A). Quercetin failed to modify neuronal viability when applied alone up to 150 μM concentration (B). Oxidative injury of P19 neurons was induced by exposure to 1.5 mM H_2_O_2_. Cell death of damaged neurons was analysed in the presence of various concentrations of quercetin by MTT assay (C) or trypan blue exclusion assay (D). Data are expressed as means ± SEM from four independent experiments. #P < 0.0001vs. cont; *P < 0.05, **P < 0.001 and ***P < 0.0001vs. H_2_O_2_-only treated group (ONE-way ANOVA followed by Tukey’s or Dunnett’s multiple comparison test).

### Quercetin did not improve H_2_O_2_-induced decrease in glutathione (GSH) content

As previously reported, quercetin prevents H_2_O_2_-provoked upregulation of ROS production.[30] Here, we analysed levels of reduced glutathione (GSH), one of the major non-enzymatic intracellular antioxidants, as oxidative stress may alter reservoirs of intracellular antioxidant systems. Exposure to H_2_O_2_ depleted GSH to 55.7% of the vehicle-treated group, but in the presence of quercetin intracellular GSH content was not restored. Comparable results were also demonstrated in mouse embryos exposed to H_2_O_2_ and quercetin [31]. Detoxification of H_2_O_2_ depends predominantly on glutathione peroxidase in a reaction that utilizes two GSH molecules, consequently leading to GSH depletion. In addition, GSH decrease in H_2_O_2_-treated neurons may result from the direct free radical scavenging, and via formation of GSH-conjugates with various electrophilic compounds [32]. However, in P19 neurons a substitution of endogenous antioxidant with an exogenous molecule with strong antioxidative activity probably provided sufficient antioxidative defence and improved survival despite the reduced GSH content.

**Fig 2.**
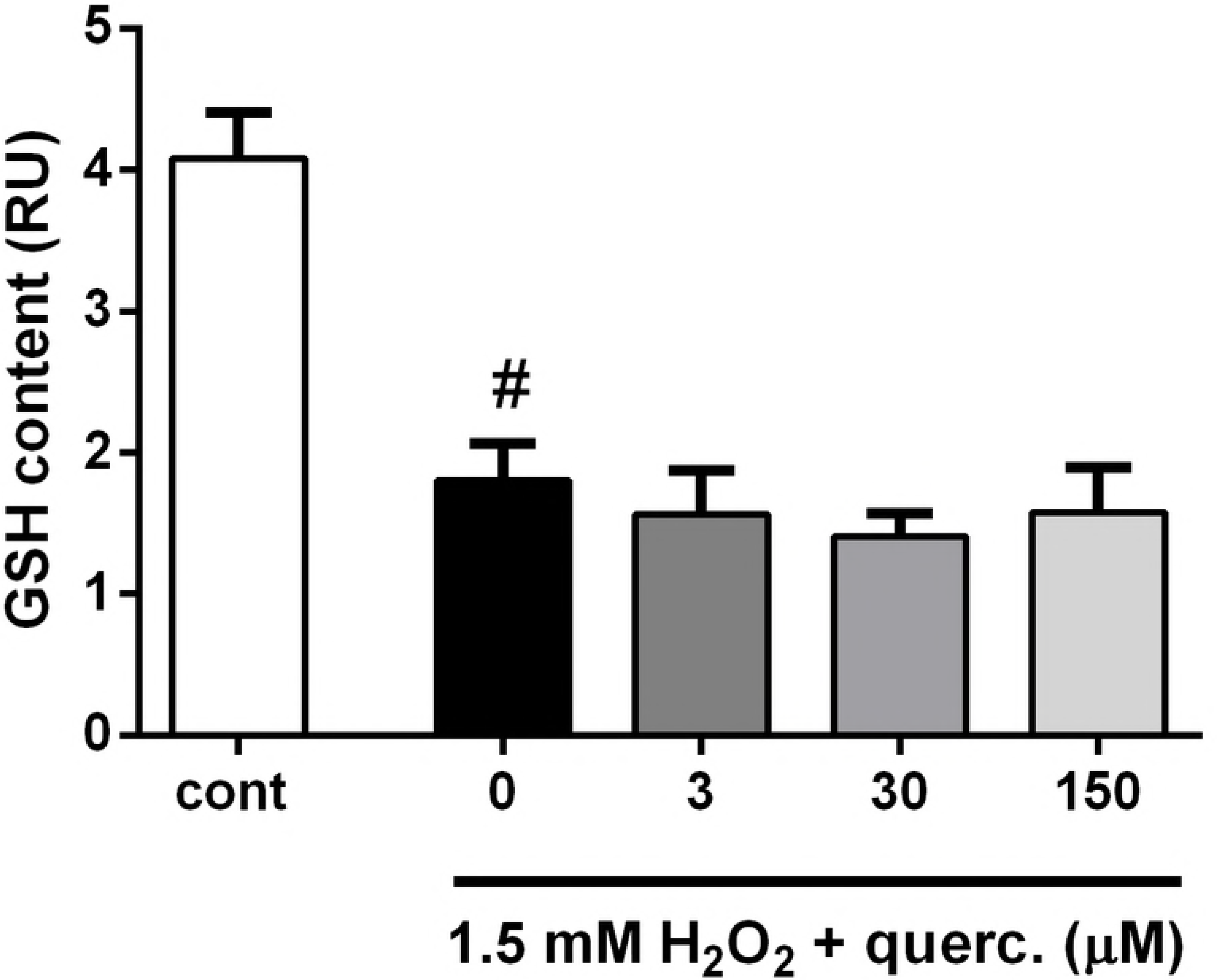
Quercetin did not affect an H_2_0_2_-induced decrease in content. At the end of 24 h treatment, GSH content was depleted in P19 neurons exposed to 1.5 mM H_2_O_2_. Presence of quercetin did not modify intracellular pool of GSH. Values represent the mean ± SEM of three independent experiments performed in triplicate. #P < 0.0001 vs. vehicle-treated group (ONE-way ANOVA followed by Tukey’s multiple comparison tests).

Neuronal death is commonly regulated by Bcl-2 family members. As represented in Figure 3A, H_2_O_2_ reduced expression of Bcl-2 mRNA. In the presence of quercetin transcriptional down-regulation of Bcl-2 was largely prevented, returning the overall Bcl-2/Bax ratio back to control values. Other studies also indicated that quercetin exerts its neuroprotective effects against oxidative stress by increasing Bcl-2 levels [33,34]. On the contrary, expression of transcription factor p53 was significantly induced by H_2_O_2_ treatment. H_2_O_2_-promoted up-regulation of p53 expression was diminished in P19 neurons simultaneously exposed to quercetin. In neuroblastoma SK-N-MC cells, the neuroprotective effect of quercetin was also accompanied by the suppression of H_2_O_2_-induced p53 enhancement [10]. Similarly, in oxidative stress induced by transient focal cerebral ischemia and reperfusion, quercetin prevented neuronal loss by diminishing p53 expression [35]. We also analysed expression of GAPDH mRNA. GAPDH is primarily viewed as a metabolic enzyme engaged in glycolysis, but it mediates numerous non-glycolytic functions, including the sensing of oxidative stress and induction of cell death [36,37]. Although H_2_O_2_ exposure did not change GAPDH expression, the pro-survival effect of quercetin correlated with pronounced GAPDH up-regulation.

**Fig 3.**
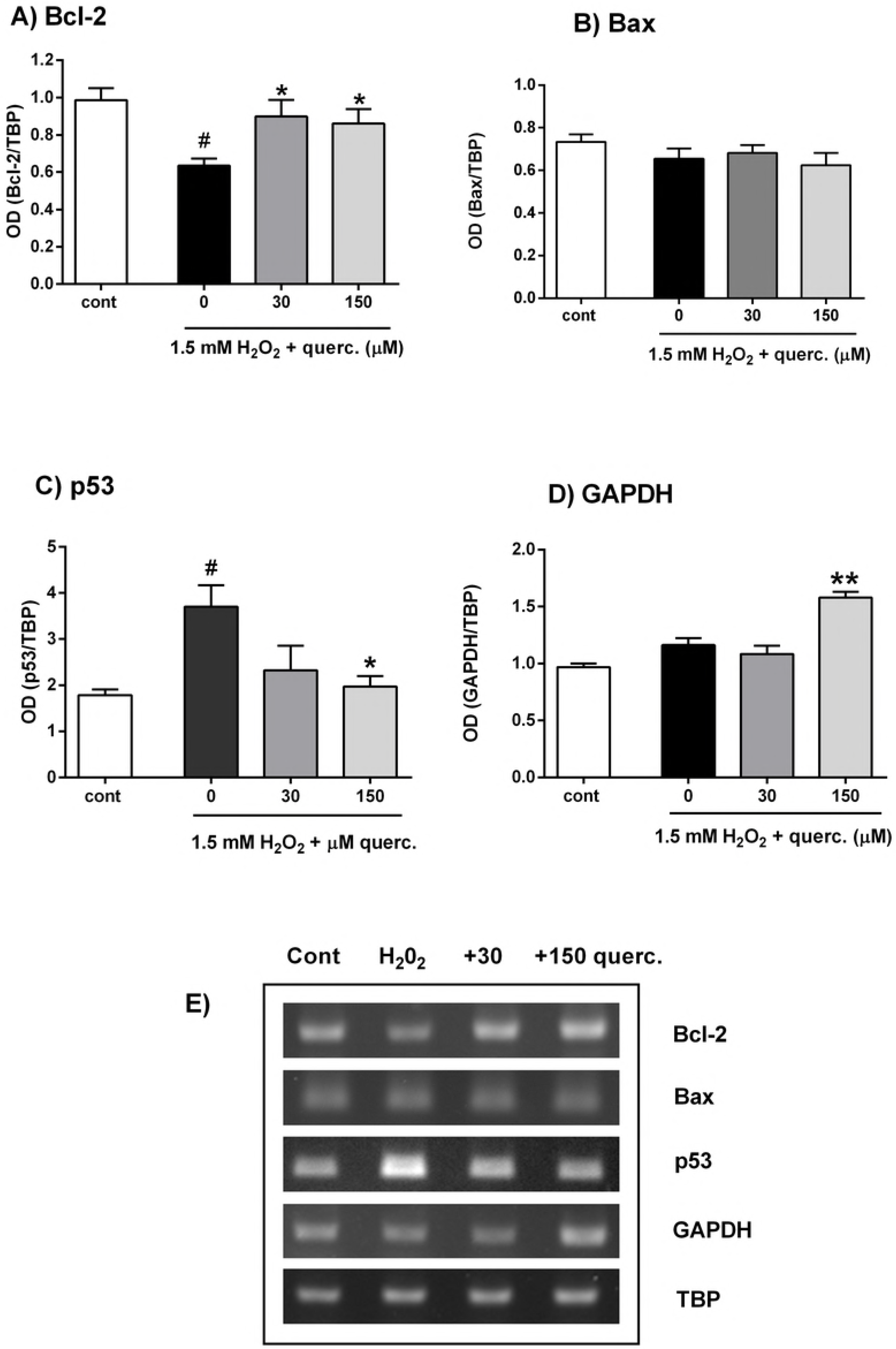
Effects of quercetin on Bcl-2, Bax, p53 and GAPDH mRNA expression during severe oxidative injury. P19 neurons were exposed to 1.5 mM H_2_O_2_ alone or in the presence of 30 and 150 μM quercetin. Total RNA was extracted and reverse transcribed into cDNA. The obtained cDNA is further amplified using specific primers. Following densitometric quantification, band intensities were normalized to the expression of housekeeping gene TBP. The data are expressed as means ± SEM from 3 independent RT-PCR analyses. #P < 0.05 vs. cont; *P < 0.05, **P < 0.01 vs. 0 group (ONE-way ANOVA and post-hoc Tukey’s multiple comparison test). Representative agarose gel electrophoresis is also shown.

H_2_O_2_ readily stimulates the conversion of sulfhydryl groups into disulfides and other oxidized species. Hence, it may inhibit or promote disulfide bonding within or between redox-sensitive cytoplasmic proteins involved in translation, glycolysis, cytoskeletal structure and antioxidative defence [38]. Following H_2_O_2_ exposure, different cysteine modifications, such as thiol oxidation and S-glutathionylation, were observed in the catalytic site of GAPDH, affecting its structure and function [37,39]. Thus, although we failed to observe H_2_O_2_-related changes in GAPDH expression, it is possible that P19 neurons have experienced a loss of at least some GAPDH functions that were restored by quercetin-induced GAPDH up-regulation. It is known that membrane-associated GAPDH binds to tubulin and regulates bundling of microtubules [36]. Hence, by stimulating microtubule bundling quercetin-mediated GAPDH up-regulation potentially could contribute to the preservation of normal neuronal branching that is disrupted by H_2_O_2_ treatment [30]. Quercetin-induced changes in the expression profile of GAPDH was also demonstrated *in vivo* [40]. The relationship between GAPDH and p53, a transcription factor that regulates expression of stress response genes is also observed. Protein p53 is upregulated and promotes death in response to diverse genotoxic cellular stresses, including oxidative stress [10,41]. In glutamate-induced oxidative stress, GAPDH translocates to the nucleus, enhances p53 expression and activates p53-mediated death pathway [42]. This suggests that quercetin-mediated decrease in p53 expression potentially could disrupt GAPDH-p53 interactions and offer neuroprotection.

P19 neurons exposed to high concentration of H_2_O_2_ died by caspase-independent apoptosis in combination with necrosis [30]. Cell death mediated by p53 may be caspase-independent [43]. The main effector of caspase-independent death program is an apoptosis-inducing factor (AIF). Overexpression of Bcl-2 (as we found in P19 neurons exposed to quercetin) may prevent AIF release and consequent cell death [44]. Necrosis also may be prevented by Bcl-2 overexpression [45]. In cultured cortical neurons interaction between death-associated protein kinase, 1 (DAPK1) and p53 was identified as a signalling point of convergence of necrosis and apoptosis [46]. They showed that activated DAPK1 phosphorylates p53. This induces expression of pro-apoptotic genes in the nucleus and initiates necrosis in the mitochondrial matrix through interactions with cyclophilin D. In addition, p53 protein can directly promote mitochondrial outer membrane permeabilization (MOMP) to trigger apoptosis [47]. Hence, by preserving the p53 and Bcl-2 expression quercetin might contribute to the survival of P19 neurons at the level of apoptotic and necrotic death events.

To investigate if the neuroprotective effects of quercetin were mediated by the modulation of intracellular signalling, P19 neurons were exposed to H_2_O_2_ and quercetin and two selected inhibitors: UO126 (an inhibitor of the Ras/Raf/MEK/ERK signalling pathway) or wortmannin (a covalent inhibitor of phosphoinositide 3-kinases (PI3K) that activates Akt/PKB). As represented in Figure 4, beneficial effects of quercetin on neuronal survival were abrogated in the presence of UO126 and wortmannin. Both inhibitors were applied in a concentration that did not affect viability when applied alone. The obtained results indicate that the neuroprotective action of quercetin was achieved through the modulation of ERK1/2 and PI3K/Akt signalling.

**Fig 4.**
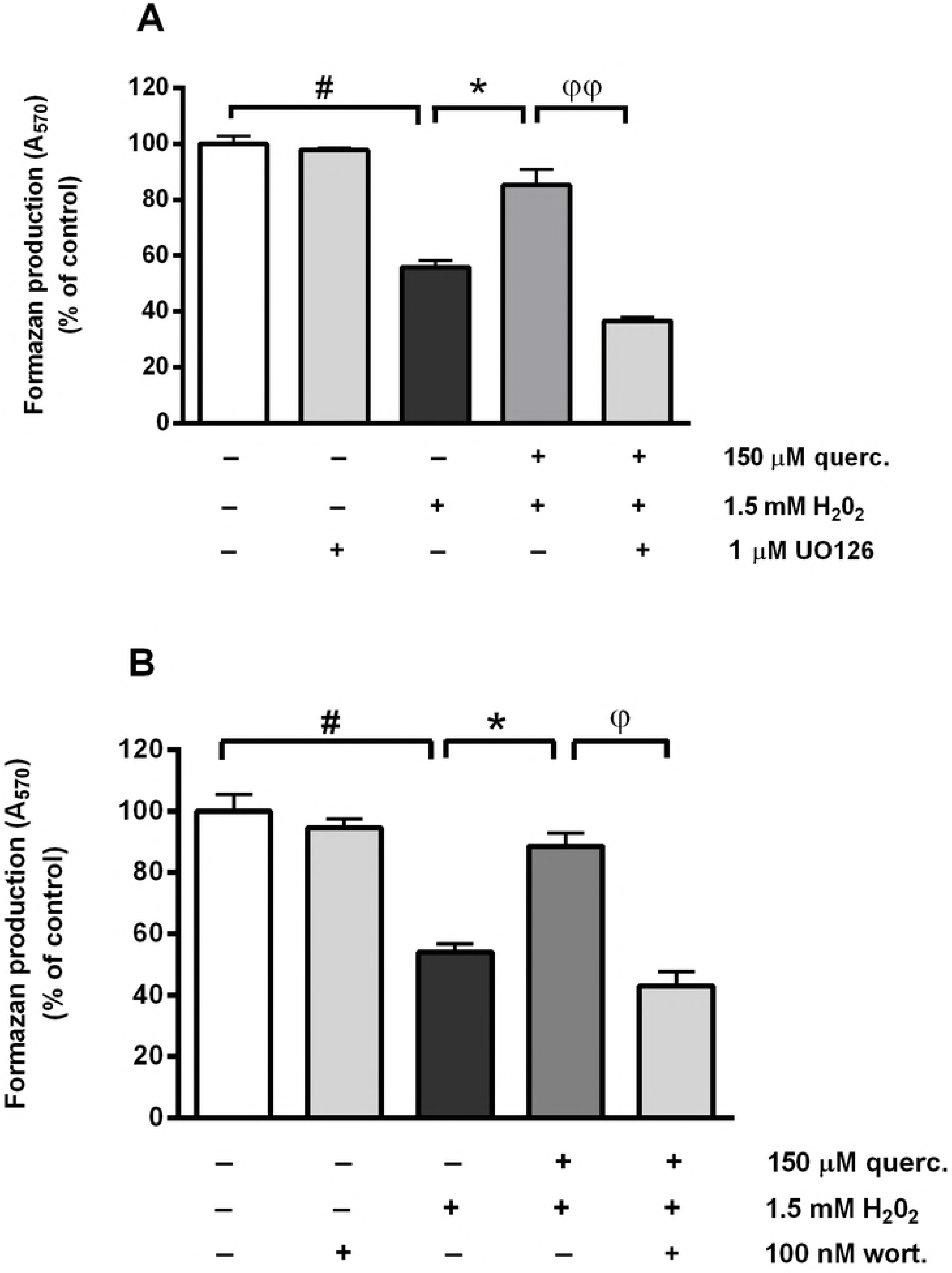
UO126 (inhibitor of the ERK1/2 pathway) and wortmannin (Akt/PKB inhibitor) prevented the neuroprotective effect of quercetin against oxidative injury. P19 neurons were treated with UO126 or wortmannin for 1 h prior to and during 24 h H_2_O_2_ treatment. UO126 applied at 1 μM concentration (upper graph), as well as 100 nM wortmannin (bottom graph) diminished survival of P19 neurons exposed simultaneously to 1.5 mM H_2_O_2_ and 150 μM quercetin for 24 h. Values are expressed as means ± SEM from three independent experiments performed in quadruplets. *P < 0.001 vs. 1 mM H_2_O_2_ alone (one-way ANOVA followed by post hoc Tukey’s test)

The PI3K/Akt and ERK1/2 signalling pathways are critically involved in controlling cell survival in response to extracellular stimuli, including H_2_O_2_-induced oxidative stress [5]. As in our study, activation of PI3K/Akt and MAPK/ERK pathways offered protection against p53-mediated cell death in sympathetic neurons.[48,49] PKB/Akt acts upstream of p53 and suppresses transcriptional activation of p53-responsive genes, [50] while signalling through both Akt and ERK1/2 pathways may be involved in the protection against H_2_O_2_-induced neuronal death through the activation of cyclic AMP regulatory-binding protein (CREB) that promotes transcription of antiapoptotic genes such as Bcl-2 [5,8]. Similarly to our results, quercetin prevented an H_2_O2-induced decrease in cell viability by inducing ERK1/2 phosphorylation, [51] and through the activation of the PI3K/Akt pathway [11]. Of note, PI3K could be involved in ERK1/2 activation in primary cortical neurons [5] and neuroblastoma cells [6]. Furthermore, quercetin may offer protection against oxidative injury by activating the Nrf2 pathway as Akt and ERK pathways participate in Nrf2 activation [9,52].

### The new biomarkers for identifying the neuroprotective effects of quercetin in H_2_O_2_-induced oxidative stress: the morphological and nanomechanical study

After we confirmed that P19 neurons have experienced the oxidative stress and that quercetin improved neuronal survival by activating the Akt and ERK pathways that normalize expression of p53 and Bax, we used AFM to image neuronal morphology and to measure membrane roughness and nanomechanical properties, parameters that have potential in biomedical applications as specific surface biomarkers for efficient detection of various cellular conditions, including oxidative stress. Although previous studies performed on live cortical neurons resulted in slightly lower Young’s moduli in comparison with values obtained on fixed P19 neurons [53,54] both types of neurons have similar elasticity maps [53,55]. This indicates that fixed P19 neurons represent a good model system to study potential biomarkers of oxidative stress and protective drugs effects at the nanoscale.

In order to more precisely identify morphological differences between treated groups, we introduced a multiple biophysical analysis by employing AFM. First, as represented in Figure 5, isolated control neurons displayed an elongated soma shape (Figure 5A), whereas neurons exposed to H_2_O_2_ have irregularly shaped and degenerated cell bodies (Figure 5C). The morphological shape of P19 neurons simultaneously treated with quercetin and H_2_O_2_ was more regular, better resembling to control neurons (Figure 5A and 5E), indicating beneficial effects of quercetin on the preservation of neuronal morphology.

**Fig 5.**
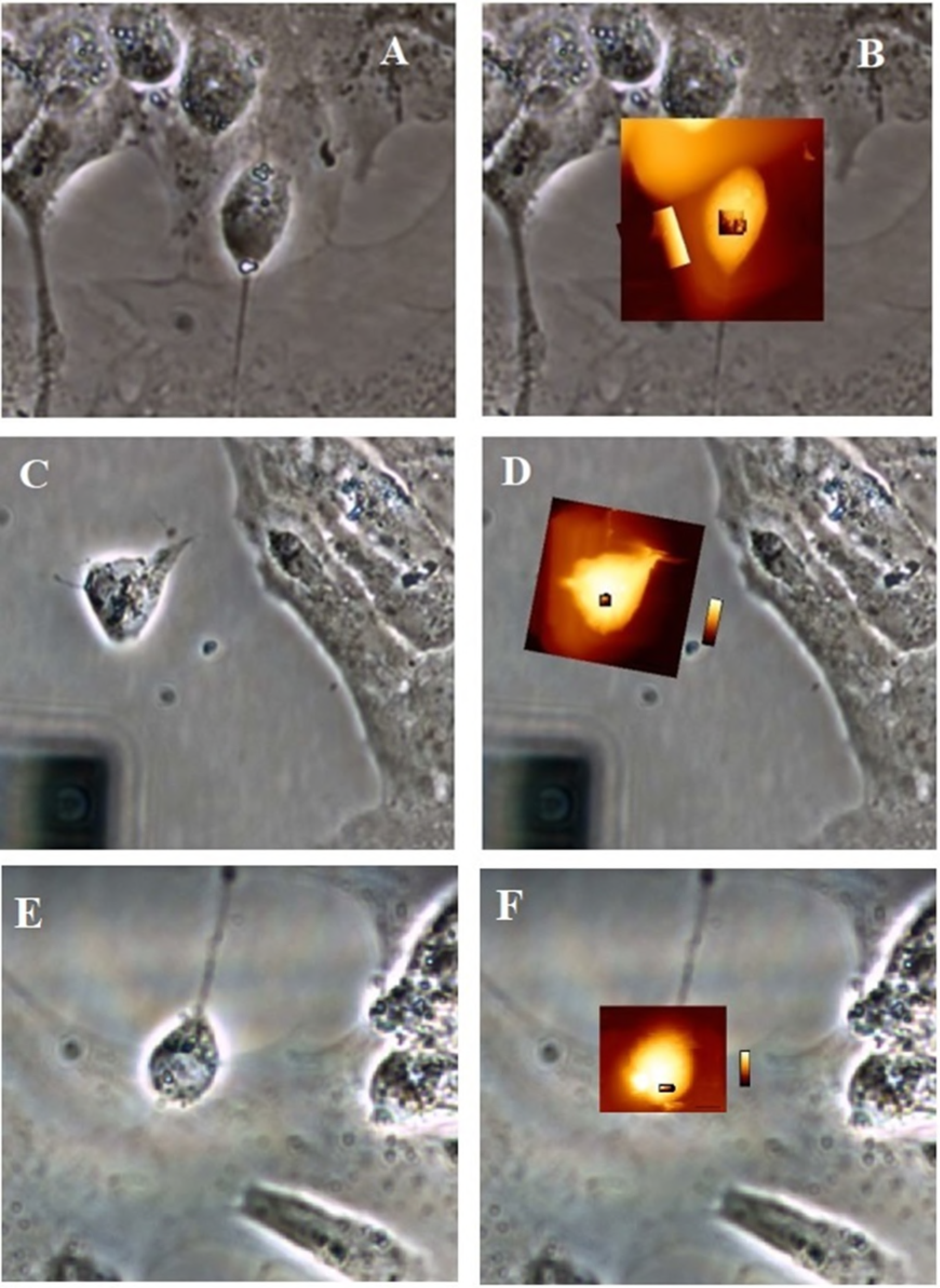
Optical images (A, C, E) and overlay of optical and QI™ images (B, D, F) of control neuronal cells (A, B), H_2_O_2_-exposed neurons (C, D) and neurons simultaneously exposed to both quercetin and H_2_O_2_ (E, F).

Second, to better reveal effects of quercetin on structural changes at the nanoscale level, we measured lateral dimensions of neuronal somas and height of the different soma regions (Figure 6) as the new potential biomarkers. As most of the analysed neurons had a somatic lateral dimension between 10 μm and 25 μm, this size range was selected for all AFM studies. The values of neuronal soma lateral dimensions (vertical *LD*_vert_ and horizontal *LD*_hor_) and height are presented in Table 2. The somatic height (h), as well as somatic lateral dimensions (LD), particularly vertical lateral dimension, LDvert of H_2_O_2_-exposed neurons, were marked lower in comparison to control. In addition, exposure to H_2_O_2_ provoked marked decrease in the cell volume, (Δ*V* = − 420 μm^3^) indicating significant rearrangement and conformational changes of cytoskeleton structure. During simultaneous treatment of neurons with H_2_O_2_ and quercetin, the observed cell volume change was approximately half of that obtained with H_2_O_2_, barely Δ*V* = - 205 μm^3^. Hence, with the applied approach using cell volume as one of the biomarkers, we were able to improve detection and recognition of protective effects of quercetin on neuronal morphology.

As evident in Figure 6B, distinct regions of control neurons contain ruffling structures probably consisting of the assembly of diverse membrane proteins and membrane folding. Similar findings of fine ruffling formations have been observed in morphological studies of neuronal growth cone [23,56]. AFM imaging revealed that individual ruffling structures are of various sizes in control neurons. These membrane protrusions were significantly suppressed in H_2_O_2_-exposed neurons (Figure 6E), probably due to the formation of higher molecular weight components by cross-linking of membrane proteins.[57] The protective effect of quercetin was particularly evident in the cross-section profile of neurons treated with both quercetin and H_2_O_2_ whose fine ruffling assemblies showed only minor modifications in comparison with control neurons (Figure 6 H).

**Fig 6.**
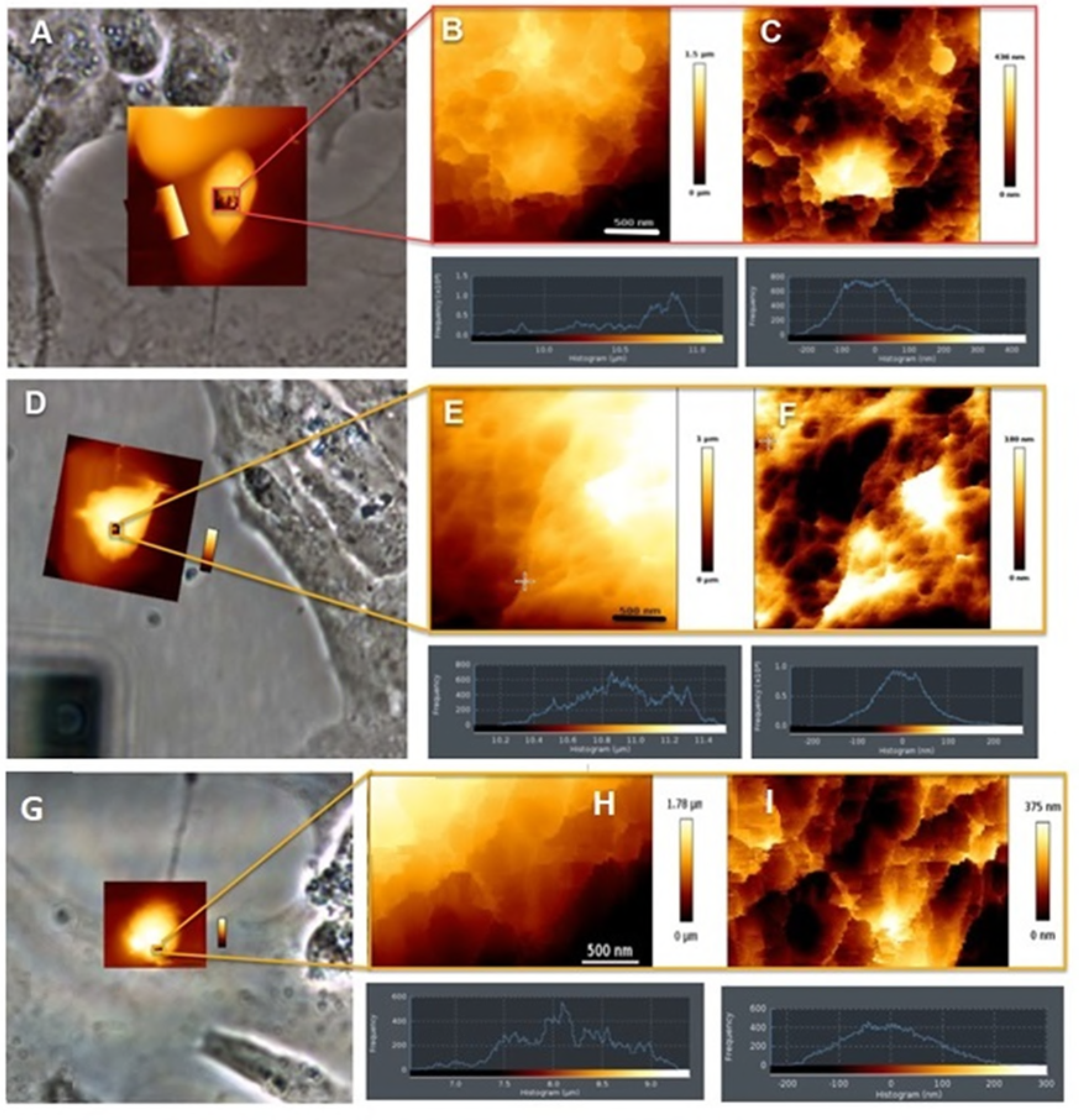
Low-resolution inverted optical microscopic images on neuron soma control system (A), on neuron after the treatment with H_2_O_2_ (D) and after the simultaneous exposure to both quercetin and H_2_O_2_ (G). The highest region of the soma was zoomed (B, E, H). The relative height differences between individual regions were consistent with data acquired from inverted optical microscopic images and indicated within histograms shown in (C, F, I). Scales are indicated. Frequency histograms of unfiltered normalized cut-off height (B, E, H bottom) and filtered normalized cut-off height (C, F, I bottom) for the line profile of a neuron.

We also performed surface roughness analysis of the neuronal soma membrane to specifically evaluate changes in membrane surface topography. A membrane roughness is an important parameter in cell studies, also with great potential for medicinal applications as a valuable biomarker. It indicates the deviation of the membrane surface topography from the ideally smooth surface. The roughness parameters described by *R*_a_ and *R*_q_ were derived from the raw height images and the height images treated with a quadratic plane fit. Ten neurons were analysed by randomly measuring several areas of each cell surface. The analysed regions of control cells were relatively small (2×2 μm^2^), but they appeared cone-shaped in cross-section profile, and their surfaces were relatively rough as shown in Table 2. The plane fit was used to subtract the curvature of the neuronal body which would otherwise influence the roughness data (Table 1). The plane fit more accurately revealed surface roughness. The scale-independent roughness parameters, obtained by computing the *R*_a_ value on a filtered profile are comparable with the dimension of protein complexes and also demonstrated membrane damage processes. While spherical crater edges in the control P19 neurons are very sharp and distinct (Figure 6B), in quercetin/H_2_O_2_ treated neurons they are pitted and became shallow (Figure 6H), and completely disappear in H_2_O_2_-treated neurons (Figure 6E). In the presence of H_2_O_2_, it is obvious that decrease in roughness parameters indicates the discrete cell membrane damage. By the addition of quercetin, changes of these parameters were markedly prevented, indicating only minor damage of the neuronal surface. Thus, reduced changes of roughness parameters indicate better structural preservation as a result of quercetin presence. Hence, roughness data as a valuable biomarker showed a protective effect of quercetin against H_2_O_2_-induced oxidative injury at the membrane level.

Recently, Lee et al. [58] demonstrated that Taxol-induced decrease in the roughness of neuroblastoma cells indicates an increase of membrane stiffness caused by microtubule translocation. In addition, they showed a time-lapse measurement of the decrease in membrane tension induced by a hypertonic solution that resulted in an increase of membrane roughness. On the contrary, our results showed that H_2_O_2_ induces a decrease in cell volume and therefore decrease in the membrane tension that resulted in the decrease of the roughness. Obviously, H_2_O_2_-induced oxidative stress in the highly organized cytoskeleton does not necessarily lead to the increase of membrane roughness. Therefore, membrane roughness can quantitatively describe cellular events at the nanoscale level including not only cytoskeletal alterations [59] and apoptotic processes [60] but also the organization of membrane components [61]. Due to the aforementioned formation of the higher molecular weight components by the crosslinking of membrane proteins, [57] the surface of H_2_O_2_-damaged membranes became markedly smoother (Figure 6F) in comparison to the rough surface of control neurons (Figure 6C). Membranes of neurons treated with both H_2_O_2_ and quercetin did not alter significantly, remaining their surface still rough (Figure 6H). Table 1 summarizes the roughness, height and lateral dimension values determined from the AFM height measurements on the highest domain of the neurons.

**Table 2.** **Roughness, height and lateral dimension data from the histograms of the neuronal detail images below**

| | $LD_{vert}/\mu m$ | $LD_{hor}/\mu m$ | $h/\mu m$ | $V/\mu m^3$ | Average $R_a/nm$ | $R_q/nm$ | Average $R_a/nm$ (quadratic plane fit) | $R_q/nm$ (quadratic plane fit) |
| --- | --- | --- | --- | --- | --- | --- | --- | --- |
| Control | 13.2±0.6 | 22.6±0.7 | 6.98±0.5 | 2080 | 280±6 | 351±7 | 77.5±0.9 | 99.1±0.4 |
| $H_2O_2$ | 17.0±0.3 | 17.1±0.9 | 5.6±1.2 | 1660 | 206±3 | 267±5 | 49.0±0.5 | 64.1±0.1 |
| Quercetin/<br>$H_2O_2$ | 17.5±0.6 | 16.5±0.4 | 6.5±0.2 | 1875 | 275±7 | 335±9 | 69.0±0.4 | 85.8±0.7 |

We further used AFM to analyse local mechanical properties of specific somatic regions by performing nanomechanical measurements. AFM provides quantitative data of the mechanical properties, as well as the direct relationship between mechanical and structural characteristics of neuronal cells [24]. Force-distance curves were acquired to determine the elastic moduli (Young’s moduli) of the different regions (Figure 7). Hertz’s model was used to fit the portions of the force curve and the resulting Young’s moduli are summarized in Table 3. As presented in the frequency histogram, significant variations in the elastic (Young’s) modulus were found between treated groups (Figures 7 B, E, H, bottom). The Young’s modulus distribution of the control neuronal soma (within the color-coded frame lines) varied between 0.5 kPa and 5 kPa with a maximum at 2.4±0.2 kPa. After H_2_O_2_ treatment it was spread at a much broader region from 0 kPa to 30 kPa and shifted its maximum towards higher value (14.7±0.5 kPa). The Young’s modulus distribution of the neurons concomitantly treated with quercetin and H_2_O_2_ varied at much narrow range (from 0 kPa to 15 kPa) and returned its maximum towards lower values (8.76±0.7 kPa). The same trend of membrane elasticity changes was observed from the detailed image of neurons, as shown in Table 3. In general, cell elasticity described by the Young’s moduli could be considered as a measure to evaluate cell integrity and physiological state. In P19 neurons changes of the Young’s modulus values obtained from the different regions were not identical. When proteins are homogeneously distributed throughout the cytoplasm and the membrane, all regions of the neuronal soma possess the similar elasticity. However, the Young’s modulus values, calculated from the overviewed and detailed images were slightly different. In Figure 7 is shown the Young modulus map described by the colour scale from dark brown (lower Young modulus value) to white colour (Higher Young modulus value). While the Young modulus value (elasticity) in control neurons are homogeneously distributed (brown colour within square in Figure 7B), the addition of H_2_O_2_ induced structural and mechanical changes that are reflected in the increased Young modulus value i.e. increased stiffness (white area within square in Figure 7E). The protective role of quercetin was observed in the discrete increase in the neuron stiffness (left side on the bottom square in Figure 7H) and confirmed that Young modulus i.e. elasticity may serve as a very sensitive biomarker for the evaluation of the level of the neuron damage. Hence, the obtained results indicate that the neuronal soma of neuronal cells contains heterogeneously arranged protein regions. H_2_O_2_-induced changes in Young’s moduli are probably determined not only with the local changes of the membrane structure but also with the local changes of the underlying cytoskeletal structure. Figures 7E, F and I show some typical frequency elastic moduli histograms taken in the highest region of the neuron. The cross-linking of membrane proteins during oxidative stress might hinder the conformational change of membrane proteins and the conformational change of microfilaments within the cytoplasm [57]. Both might be the reason for the increased membrane Young’s modulus i.e. increased stiffness upon H_2_O_2_ treatment. In comparison to H_2_O_2_ exposure, a reduced increase in stiffness was observed during simultaneous exposure to both quercetin and H_2_O_2_. These results suggest that quercetin offers protection towards certain mechano-chemical processes induced by oxidative environment that affect cytoskeleton and membrane organization in the investigated regions of the neuronal soma.

**Fig 7.**
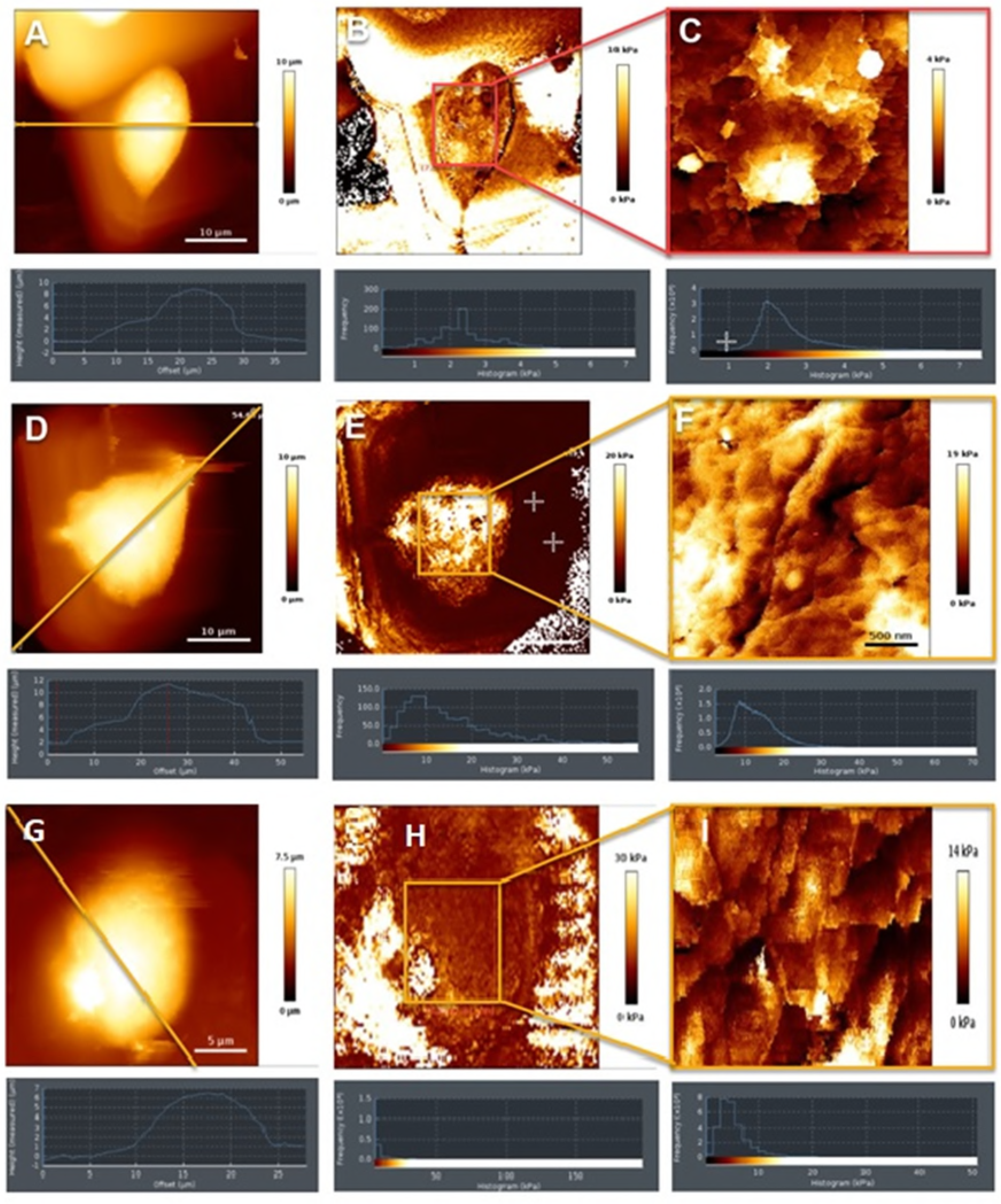
The height topographic image of specific somatic regions in control system (A), H_2_O_2_-exposed neurons (D) and in neurons simultaneously exposed to both quercetin and H_2_O_2_ (G) determined by nanomechanical measurements using AFM. The nanomechanical mapping and histogram of Young’ modulus of the specific somatic region in control system (B), H_2_O_2_-exposed neurons (E) and neurons simultaneously exposed to both quercetin and H_2_O_2_ (H). Colour-coded frame lines identified zoomed domain of control system (C) H_2_O_2_-exposed neurons (F) and neurons simultaneously exposed to both quercetin and H_2_O_2_ (I).

**Table 3.**
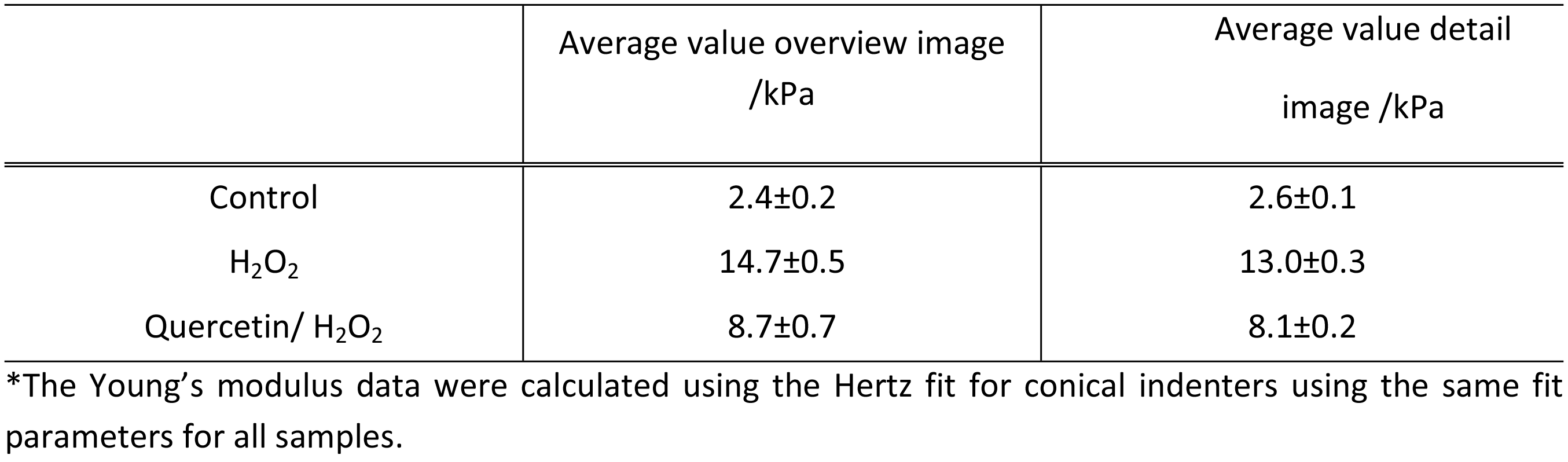
Summarizing Young’s modulus data (average values from the histograms)

The presented data performed on fixed P19 neurons provides detailed morphological changes of cellular structures under different treatment regimes. First, we demonstrated inner structural reorganization of the cytoskeleton in oxidative conditions, manifested by increased cell somatic lateral dimensions, height and cell volume. The observed alterations in morphology were suppressed by quercetin presence confirming its neuroprotective role. Second, at the membrane level, the observed morphological changes were reflected in distinct roughness values of treated neurons. The membrane roughness which originates from individual short and long protein ruffling structures [23] was decreased in P19 neurons exposed to H_2_O_2_, but turned back towards initial state during simultaneous H_2_O_2_ and quercetin treatment. H_2_O_2_ probably induces conformational changes of actin structures and promotes degradation of highly oriented bundles, consequently leading to their shortening which corresponds to the lower values of roughness parameters. Third, AFM was used to perform nanomechanical measurements in specific regions of P19 neurons. We observed that stiffer somatic regions simultaneously exhibited lower elasticity, i.e. higher Young’s modulus value. The stiffness (considered as the absence of the cell elasticity) of H_2_0_2_-treated neurons was markedly increased and then reduced almost to half in the presence of quercetin. Finally, we observed a strong correlation between cell stiffness and roughness parameters during H_2_O_2_-induced injury.

As a new biophysical approach in the study of oxidative stress-induced neurodegeneration, we used AFM for revealing protective effects of quercetin at the nanoscale resolution. The obtained results suggest that AFM and QI™ mode are powerful tools to characterize the morphology and resolve stiffness differences of the neuronal cell bodies. If combined with standard cellular and molecular methods, AFM can greatly improve detection and recognition of effects of various drugs and toxins on cellular morphology and physiology using very sensitive biomarkers as roughness and elasticity. In a wider context, a better understanding of these effects can be used for drug screening, and potentially to the development of novel therapeutic strategies. This AFM study represents the first detailed analysis of the protective effects of quercetin on neuronal membrane and cytoskeleton organization and in general, indicates a great potential of AFM in biomedical research. If combined with standard cellular and molecular methods, AFM can greatly improve detection and recognition of effects of various drugs and toxins on cellular morphology and physiology.

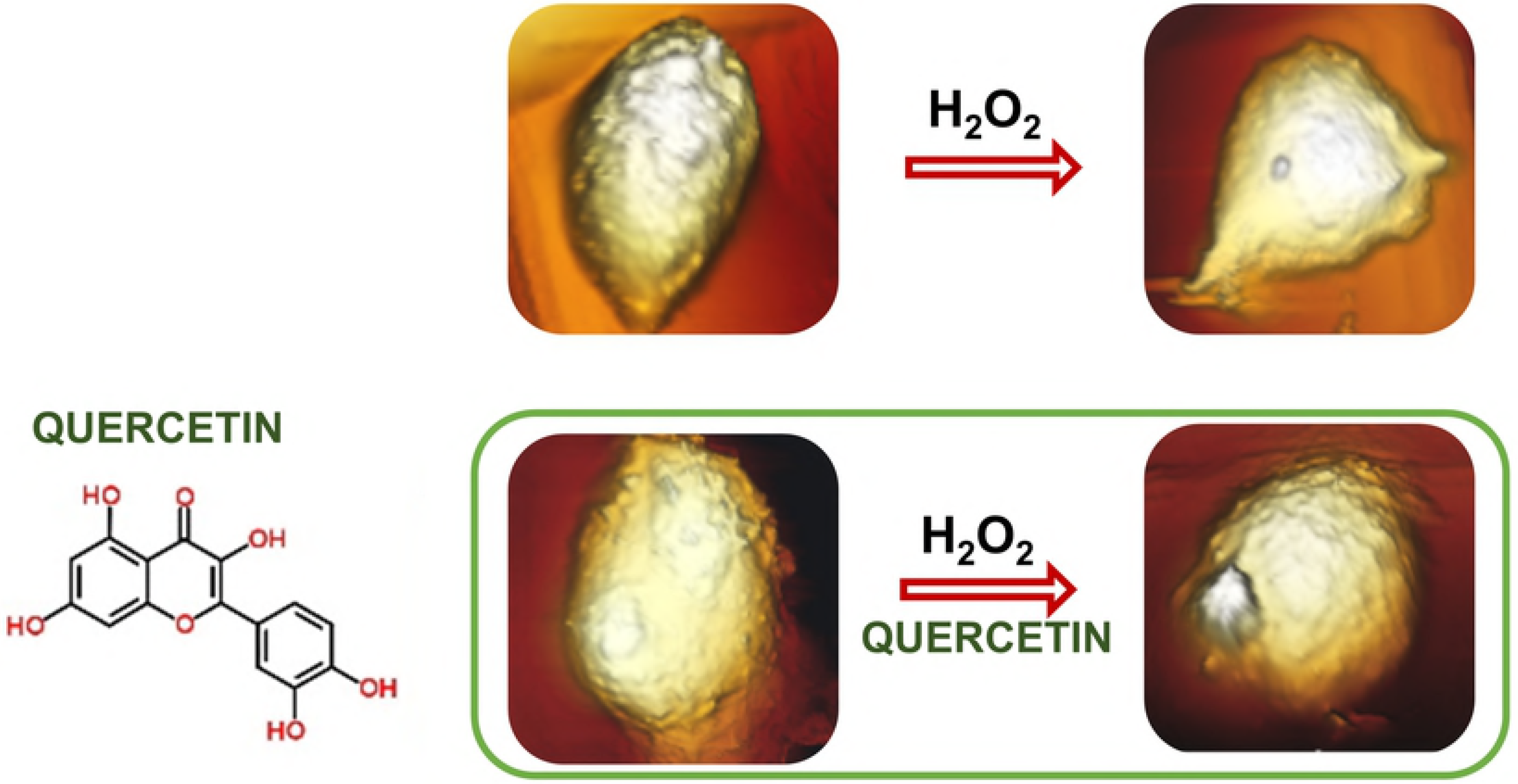

## Acknowledgement

This work was supported by Croatian Science Foundation [grant ref. IP-2016-06-8415]. The authors thank the JPK Instruments AG, Berlin, Germany for the help within the AFM measurements and analyses.

## Authors contribution

S. Šegota and M. Jazvinšćak Jembrek: critically analysed the data and wrote corresponding parts of the text; V. Čadeš and S. Šegota performed the AFM measurement and data analysis; M. Javinšćak Jembrek and J. Vlainic: performed the P19 cell culturing and P19 neuronal differentiation, drug treatment, Assessment of cell death and all biological assays; S. Šegota, M. Jazvinšćak Jembrek, V. Čadež and J. Vlainić contributed to the interpretation of the results.

## References

1. Gandhi S, Abramov AY. Mechanism of oxidative stress in neurodegeneration. Oxid Med Cell Longev. 2012;2012: 428010. doi:10.1155/2012/428010

2. Wang X, Michaelis EK. Selective neuronal vulnerability to oxidative stress in the brain. Front Aging Neurosci. 2010;2: 12. doi:10.3389/fnagi.2010.00012

3. Bienert GP, Schjoerring JK, Jahn TP. Membrane transport of hydrogen peroxide. Biochim Biophys Acta. 2006;1758: 994–1003. doi:10.1016/j.bbamem.2006.02.015

4. Lennicke C, Rahn J, Lichtenfels R, Wessjohann LA, Seliger B. Hydrogen peroxide - production, fate and role in redox signaling of tumor cells. Cell Commun Signal CCS. 2015;13: 39. doi:10.1186/s12964-015-0118-6

5. Crossthwaite AJ, Hasan S, Williams RJ. Hydrogen peroxide-mediated phosphorylation of ERK1/2, Akt/PKB and JNK in cortical neurones: dependence on Ca(2+) and PI3-kinase. J Neurochem. 2002;80: 24–35.

6. Ruffels J, Griffin M, Dickenson JM. Activation of ERK1/2, JNK and PKB by hydrogen peroxide in human SH-SY5Y neuroblastoma cells: role of ERK1/2 in H_2_O_2_-induced cell death. Eur J Pharmacol. 2004;483: 163–173. doi:10.1016/j.ejphar.2003.10.032

7. Wang X, Michaelis EK. Selective neuronal vulnerability to oxidative stress in the brain. Front Aging Neurosci. 2010;2: 12. doi:10.3389/fnagi.2010.00012

8. Dajas F. Life or death: neuroprotective and anticancer effects of quercetin. J Ethnopharmacol. 2012;143: 383–396. doi:10.1016/j.jep.2012.07.005

9. Costa LG, Garrick JM, Roquè PJ, Pellacani C. Mechanisms of Neuroprotection by Quercetin: Counteracting Oxidative Stress and More. Oxid Med Cell Longev. 2016;2016: 2986796. doi:10.1155/2016/2986796

10. Roshanzamir F, Yazdanparast R. Quercetin attenuates cell apoptosis of oxidant-stressed SK-N-MC cells while suppressing up-regulation of the defensive element, HIF-1α. Neuroscience. 2014;277: 780–793. doi:10.1016/j.neuroscience.2014.07.036

11. Chen L, Sun L, Liu Z, Wang H, Xu C. Protection afforded by quercetin against H_2_O_2_-induced apoptosis on PC12 cells via activating PI3K/Akt signal pathway. J Recept Signal Transduct Res. 2016;36: 98–102. doi:10.3109/10799893.2015.1049363

12. Spencer JPE. The interactions of flavonoids within neuronal signalling pathways. Genes Nutr. 2007;2: 257–273. doi:10.1007/s12263-007-0056-z

13. Kelsey NA, Wilkins HM, Linseman DA. Nutraceutical antioxidants as novel neuroprotective agents. Mol Basel Switz. 2010;15: 7792–7814. doi:10.3390/molecules15117792

14. Echeverry C, Arredondo F, Abin-Carriquiry JA, Midiwo JO, Ochieng C, Kerubo L, et al. Pretreatment with natural flavones and neuronal cell survival after oxidative stress: a structure-activity relationship study. J Agric Food Chem. 2010;58: 2111–2115. doi:10.1021/jf902951v

15. Notas G, Nifli A-P, Kampa M, Pelekanou V, Alexaki V-I, Theodoropoulos P, et al. Quercetin accumulates in nuclear structures and triggers specific gene expression in epithelial cells. J Nutr Biochem. 2012;23: 656–666. doi:10.1016/j.jnutbio.2011.03.010

16. Arredondo F, Echeverry C, Abin-Carriquiry JA, Blasina F, Antúnez K, Jones DP, et al. After cellular internalization, quercetin causes Nrf2 nuclear translocation, increases glutathione levels, and prevents neuronal death against an oxidative insult. Free Radic Biol Med. 2010;49: 738–747. doi:10.1016/j.freeradbiomed.2010.05.020

17. Ishikawa Y, Kitamura M. Anti-apoptotic effect of quercetin: intervention in the JNK-and ERK-mediated apoptotic pathways. Kidney Int. 2000;58: 1078–1087. doi:10.1046/j.1523-1755.2000.00265.x

18. Baptista FI, Henriques AG, Silva AMS, Wiltfang J, da Cruz e Silva OAB. Flavonoids as therapeutic compounds targeting key proteins involved in Alzheimer’s disease. ACS Chem Neurosci. 2014;5: 83–92. doi:10.1021/cn400213r

19. Youl E, Bardy G, Magous R, Cros G, Sejalon F, Virsolvy A, et al. Quercetin potentiates insulin secretion and protects INS-1 pancreatic β-cells against oxidative damage via the ERK1/2 pathway. Br J Pharmacol. 2010;161: 799–814. doi:10.1111/j.1476-5381.2010.00910.x

20. Li M. Introduction to Atomic Force Microscopy-Based Nanorobotics for Biomedical Applications. Investigations of Cellular and Molecular Biophysical Properties by Atomic Force Microscopy Nanorobotics. Springer, Singapore; 2018. pp. 1–20. doi:10.1007/978-981-10-6829-4_1

21. Mogilner A, Keren K. The shape of motile cells. Curr Biol CB. 2009;19: R762–771. doi:10.1016/j.cub.2009.06.053

22. Fletcher DA, Mullins RD. Cell mechanics and the cytoskeleton. Nature. 2010;463: 485–492. doi:10.1038/nature08908

23. Xiong Y, Lee AC, Suter DM, Lee GU. Topography and Nanomechanics of Live Neuronal Growth Cones Analyzed by Atomic Force Microscopy. Biophys J. 2009;96: 5060–5072. doi:10.1016/j.bpj.2009.03.032

24. Jembrek MJ, Šimić G, Hof PR, Šegota S. Atomic force microscopy as an advanced tool in neuroscience. Transl Neurosci. 2015;6: 117–130. doi:10.1515/tnsci-2015-0011

25. Spedden E, Staii C. Neuron Biomechanics Probed by Atomic Force Microscopy. Int J Mol Sci. 2013;14: 16124–16140. doi:10.3390/ijms140816124

26. D’Agostino DP, Olson JE, Dean JB. Acute hyperoxia increases lipid peroxidation and induces plasma membrane blebbing in human U87 glioblastoma cells. Neuroscience. 2009;159: 1011–1022. doi:10.1016/j.neuroscience.2009.01.062

27. Xilouri M, Papazafiri P. Anti-apoptotic effects of allopregnanolone on P19 neurons. Eur J Neurosci. 2006;23: 43–54. doi:10.1111/j.1460-9568.2005.04548.x

28. Jembrek MJ, Gašparović AČ, Vuković L, Vlainić J, Žarković N, Oršolić N. Quercetin supplementation: insight into the potentially harmful outcomes of neurodegenerative prevention. Naunyn Schmiedebergs Arch Pharmacol. 2012;385: 1185–1197. doi:10.1007/s00210-012-0799-y

29. Sader JE, Chon JWM, Mulvaney P. Calibration of rectangular atomic force microscope cantilevers. Rev Sci Instrum. 1999;70: 3967–3969. doi:10.1063/1.1150021

30. Jazvinšćak Jembrek M, Vuković L, Puhović J, Erhardt J, Oršolić N. Neuroprotective effect of quercetin against hydrogen peroxide-induced oxidative injury in P19 neurons. J Mol Neurosci MN. 2012;47: 286–299. doi:10.1007/s12031-012-9737-1

31. Yu S, Long H, Lyu Q, Zhang Q, Yan Z, Liang H, et al. Protective effect of quercetin on the development of preimplantation mouse embryos against hydrogen peroxide-induced oxidative injury. PloS One. 2014;9: e89520. doi:10.1371/journal.pone.0089520

32. Dickinson DA, Forman HJ. Cellular glutathione and thiols metabolism. Biochem Pharmacol. 2002;64: 1019–1026.

33. Suematsu N, Hosoda M, Fujimori K. Protective effects of quercetin against hydrogen peroxide-induced apoptosis in human neuronal SH-SY5Y cells. Neurosci Lett. 2011;504: 223–227. doi:10.1016/j.neulet.2011.09.028

34. Sharma DR, Wani WY, Sunkaria A, Kandimalla RJ, Sharma RK, Verma D, et al. Quercetin attenuates neuronal death against aluminum-induced neurodegeneration in the rat hippocampus. Neuroscience. 2016;324: 163–176. doi:10.1016/j.neuroscience.2016.02.055

35. Ahmad A, Khan MM, Hoda MN, Raza SS, Khan MB, Javed H, et al. Quercetin protects against oxidative stress associated damages in a rat model of transient focal cerebral ischemia and reperfusion. Neurochem Res. 2011;36: 1360–1371. doi:10.1007/s11064-011-0458-6

36. Sirover MA. New insights into an old protein: the functional diversity of mammalian glyceraldehyde-3-phosphate dehydrogenase. Biochim Biophys Acta. 1999;1432: 159–184.

37. Butterfield DA, Hardas SS, Lange MLB. Oxidatively modified glyceraldehyde-3-phosphate dehydrogenase (GAPDH) and Alzheimer’s disease: many pathways to neurodegeneration. J Alzheimers Dis JAD. 2010;20: 369–393. doi:10.3233/JAD-2010-1375

38. Cumming RC, Andon NL, Haynes PA, Park M, Fischer WH, Schubert D. Protein disulfide bond formation in the cytoplasm during oxidative stress. J Biol Chem. 2004;279: 21749–21758. doi:10.1074/jbc.M312267200

39. Hwang NR, Yim S-H, Kim YM, Jeong J, Song EJ, Lee Y, et al. Oxidative modifications of glyceraldehyde-3-phosphate dehydrogenase play a key role in its multiple cellular functions. Biochem J. 2009;423: 253–264. doi:10.1042/BJ20090854

40. Natsume Y, Kadota K, Satsu H, Shimizu M. Effect of Quercetin on the Gene Expression Profile of the Mouse Intestine. Biosci Biotechnol Biochem. 2009;73: 722–725. doi:10.1271/bbb.80484

41. Steckley D, Karajgikar M, Dale LB, Fuerth B, Swan P, Drummond-Main C, et al. Puma is a dominant regulator of oxidative stress induced Bax activation and neuronal apoptosis. J Neurosci Off J Soc Neurosci. 2007;27: 12989–12999. doi:10.1523/JNEUROSCI.3400-07.2007

42. Zhai D, Chin K, Wang M, Liu F. Disruption of the nuclear p53-GAPDH complex protects against ischemia-induced neuronal damage. Mol Brain. 2014;7: 20. doi:10.1186/1756-6606-7-20

43. Johnson MD, Kinoshita Y, Xiang H, Ghatan S, Morrison RS. Contribution of p53-dependent caspase activation to neuronal cell death declines with neuronal maturation. J Neurosci Off J Soc Neurosci. 1999;19: 2996–3006.

44. Cregan SP, Dawson VL, Slack RS. Role of AIF in caspase-dependent and caspase-independent cell death. Oncogene. 2004;23: 2785–2796. doi:10.1038/sj.onc.1207517

45. Kane DJ, Ord T, Anton R, Bredesen DE. Expression of bcl-2 inhibits necrotic neural cell death. J Neurosci Res. 1995;40: 269–275. doi:10.1002/jnr.490400216

46. Pei L, Shang Y, Jin H, Wang S, Wei N, Yan H, et al. DAPK1-p53 interaction converges necrotic and apoptotic pathways of ischemic neuronal death. J Neurosci Off J Soc Neurosci. 2014;34: 6546–6556. doi:10.1523/JNEUROSCI.5119-13.2014

47. Nikoletopoulou V, Markaki M, Palikaras K, Tavernarakis N. Crosstalk between apoptosis, necrosis and autophagy. Biochim Biophys Acta. 2013;1833: 3448–3459. doi:10.1016/j.bbamcr.2013.06.001

48. Philpott KL, McCarthy MJ, Klippel A, Rubin LL. Activated phosphatidylinositol 3-kinase and Akt kinase promote survival of superior cervical neurons. J Cell Biol. 1997;139: 809–815.

49. Anderson CN, Tolkovsky AM. A role for MAPK/ERK in sympathetic neuron survival: protection against a p53-dependent, JNK-independent induction of apoptosis by cytosine arabinoside. J Neurosci Off J Soc Neurosci. 1999;19: 664–673.

50. Yamaguchi A, Tamatani M, Matsuzaki H, Namikawa K, Kiyama H, Vitek MP, et al. Akt activation protects hippocampal neurons from apoptosis by inhibiting transcriptional activity of p53. J Biol Chem. 2001;276: 5256–5264. doi:10.1074/jbc.M008552200

51. Youl E, Bardy G, Magous R, Cros G, Sejalon F, Virsolvy A, et al. Quercetin potentiates insulin secretion and protects INS-1 pancreatic β-cells against oxidative damage via the ERK1/2 pathway. Br J Pharmacol. 2010;161: 799–814. doi:10.1111/j.1476-5381.2010.00910.x

52. Wang Q, Chuikov S, Taitano S, Wu Q, Rastogi A, Tuck SJ, et al. Dimethyl Fumarate Protects Neural Stem/Progenitor Cells and Neurons from Oxidative Damage through Nrf2-ERK1/2 MAPK Pathway. Int J Mol Sci. 2015;16: 13885–13907. doi:10.3390/ijms160613885

53. Spedden E, White JD, Naumova EN, Kaplan DL, Staii C. Elasticity Maps of Living Neurons Measured by Combined Fluorescence and Atomic Force Microscopy. Biophys J. 2012;103: 868–877. doi:10.1016/j.bpj.2012.08.005

54. Spedden E, White JD, Kaplan D, Staii C. Young’s Modulus of Cortical and P19 Derived Neurons Measured by Atomic Force Microscopy. MRS Online Proc Libr Arch. 2012;1420. doi:10.1557/opl.2012.485

55. Bain G, Ray WJ, Yao M, Gottlieb DI. From embryonal carcinoma cells to neurons: the P19 pathway. BioEssays News Rev Mol Cell Dev Biol. 1994;16: 343–348. doi:10.1002/bies.950160509

56. Rochlin MW, Dailey ME, Bridgman PC. Polymerizing Microtubules Activate Site-directed F-Actin Assembly in Nerve Growth Cones. Mol Biol Cell. 1999;10: 2309–2327.

57. Wang X, Wu Z, Song G, Wang H, Long M, Cai S. Effects of oxidative damage of membrane protein thiol groups on erythrocyte membrane viscoelasticities. Clin Hemorheol Microcirc. 1999;21: 137–146.

58. Lee, C. W., Jang, L. L., Pan, H. J., Chen, Y. R., Chen, C. C., Lee, C. H., 2016, Membrane roughness as a sensitive parameter reflecting the status of neuronal cells in response to chemical and nanoparticle treatments. l. J Nanobiotechnol 14, 9.

59. Girasole M, Pompeo G, Cricenti A, Congiu-Castellano A, Andreola F, Serafino A, et al. Roughness of the plasma membrane as an independent morphological parameter to study RBCs: a quantitative atomic force microscopy investigation. Biochim Biophys Acta. 2007;1768: 1268–1276. doi:10.1016/j.bbamem.2007.01.014

60. Kim KS, Cho CH, Park EK, Jung M-H, Yoon K-S, Park H-K. AFM-Detected Apoptotic Changes in Morphology and Biophysical Property Caused by Paclitaxel in Ishikawa and HeLa Cells. PLOS ONE. 2012;7: e30066. doi:10.1371/journal.pone.0030066

61. Reister E, Bihr T, Seifert U, Smith A-S. Two intertwined facets of adherent membranes: membrane roughness and correlations between ligand-receptors bonds. New J Phys. 2011;13: 025003. doi:10.1088/1367-2630/13/2/025003

